# OmicsLake: Versioned, Agent-Aware Data Lineage for R/Bioconductor Workflows

**DOI:** 10.64898/2026.07.17.739088

**Authors:** Yusuke Matsui

**Affiliations:** Institute for Glyco-core Research (iGCORE), Nagoya University, Nagoya, Japan; Biomedical and Health Informatics Unit, Graduate School of Medicine, Nagoya University, Nagoya, Japan

**Keywords:** reproducibility, data provenance, data versioning, lineage tracking, agentic bioinformatics, DuckDB

## Abstract

OmicsLake is an R package for version-aware, dataset-level lineage in R/Bioconductor exploratory workflows. It associates stored versions with dependencies captured within tracked `ref()–dplyr–save_as()` pipelines or supplied explicitly, and can attach caller-supplied human or software-agent identifiers. Controlled tests recovered all ground-truth edges within the supported path and enabled explicit recovery of an overwritten tagged intermediate. OmicsLake 0.99.3 is MIT-licensed software available at https://github.com/matsui-lab/OmicsLake (tag `v0.99.3`); source, tests, and evaluation files are archived at https://doi.org/10.5281/zenodo.21391428. Supplementary Material accompanies this preprint.

## 1 Introduction

Reproducibility remains a fundamental challenge in computational biology (Peng, 2011; Sandve et al., 2013), and failures can arise from unrecorded intermediate data states as well as from code and environment. In an audit of 15,817 biomedical Jupyter notebooks, dependencies could be installed for 10,388 and 879 then reproduced identically (8.5% of the dependency-installed subset) (Samuel and Mietchen, 2024). Governance is also relevant when large language model (LLM) agents participate in analysis (Zhou et al., 2025): in a separate benchmark of 300 agent-generated software projects, 31.7% did not execute out of the box (Vangala et al., 2025). A useful provenance record should connect an analytical result to the data version used, its upstream datasets, and any supplied human or software-agent identifier. Typical omics analyses—normalization, filtering, differential expression, and enrichment—produce interdependent states that are repeatedly revised during exploration (Love et al., 2014); Bioconductor (Huber et al., 2015) standardizes containers such as SummarizedExperiment (Morgan et al., 2025), but these containers do not themselves version evolving states or record derivation lineage.

Existing tools address different parts of this problem. Workflow managers such as Snakemake (Köster and Rahmann, 2012), Nextflow (Di Tommaso et al., 2017), and targets (Landau, 2021) provide pipeline-level provenance from predefined specifications. Data catalogs such as pins (Silge et al., 2026) and DVC (Iterative, 2020) version data, while Git LFS (Git LFS contributors, 2024) operates at file granularity. DuckLake (Raasveldt and Mühleisen, 2025) provides versioned Parquet tables with snapshots and time travel through DuckDB. Among the tools evaluated here, these capabilities were not combined with automatic ad hoc derivation lineage, supplied agent identifiers, in-process queries, and native Bioconductor object handling.

OmicsLake provides version-aware dataset lineage over a queryable local store. It automatically propagates source metadata through supported dplyr verbs, including two-input joins, when a pipeline starts with `ref()` and ends with `save_as()` ; package boundaries and other untracked operations use explicit `depends_on` annotations. Stored versions can also carry prompt, run, agent, and script identifiers supplied by a calling harness. Native adapters round-trip Bioconductor S4 objects so that the versioned unit can remain the biological object rather than only a flattened table. OmicsLake complements Git (code), renv (Ushey and Wickham, 2024) (R packages/environment), and Docker (system/OS) by addressing the intermediate data-state layer.

## 2 Implementation

OmicsLake is an R6-based package combining three embedded backends (Fig. 1): DuckDB (Raasveldt and Mühleisen, 2019), an analytical database running within the R process; Apache Arrow (Apache Arrow developers, 2016), for in-memory interchange; and Parquet (Apache Parquet developers, 2013), for columnar export with configurable compression. Users install a single R package without operating an external database service. The current representation prioritizes queryability over minimal file size: in the dense numeric benchmark, Parquet occupied 35.2 MB versus 29.5 MB for compression-enabled RDS (19.3% larger).

**Figure 1.**
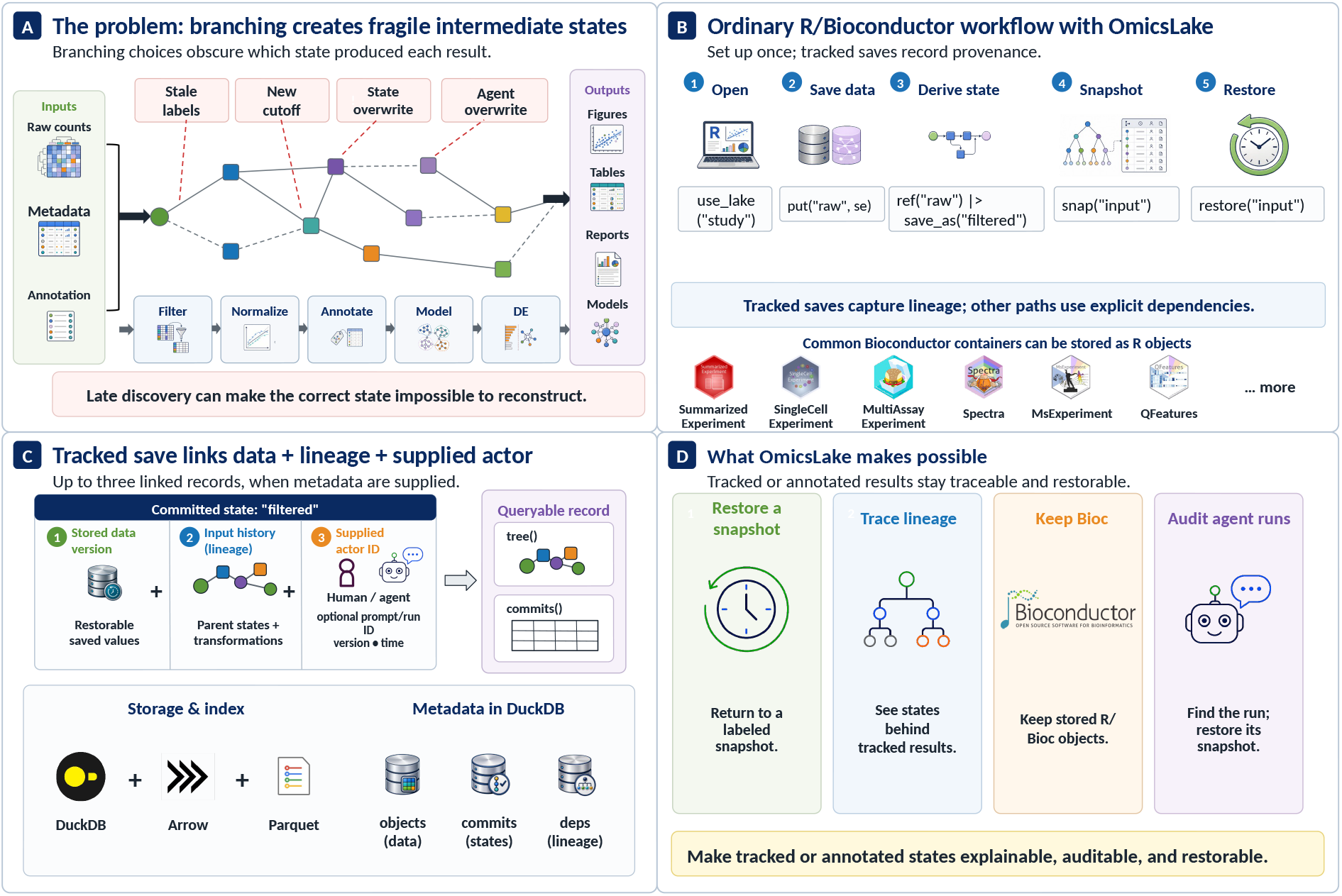
Conceptual overview of OmicsLake. (A) Branching exploratory workflows create fragile intermediate states when labels, thresholds, or human/agent runs change. (B) OmicsLake fits ordinary R/Bioconductor analysis: users open a lake, save and derive data, create a labeled snapshot, and restore that snapshot. (C) Tracked saves can associate stored versions with captured input lineage and, when supplied, human or software-agent identifiers in queryable metadata backed by DuckDB, Arrow, and Parquet. (D) The resulting record supports labeled-snapshot restoration, lineage tracing for tracked or annotated results, stored Bioconductor objects, and audit of agent-associated intermediates.

The API is a small set of orthogonal operations (Supplementary Table S2 and Figure S1): `put` /`get` auto-detect type, routing tabular data to DuckDB and arbitrary R objects to serialized storage; `ref()` returns lazy, dplyr-compatible references enabling DuckDB query pushdown; `snap` /`restore` create and roll back labeled snapshots; and `save_as()` terminates a dplyr pipe, persisting results while carrying inferred dependencies. For tracked dplyr workflows, `ref()` attaches source metadata via a `lake_tbl` S3 class. Methods for supported single-table verbs and two-input joins preserve and merge this metadata, and `save_as()` records the accumulated sources as lineage edges, from which `tree()` reconstructs the directed acyclic graph (DAG). Operations crossing package boundaries, including Bioconductor-specific functions, require `depends_on`, `with_tracking()`, or another explicit tracking context.

The distinguishing primitive is an optional *three-way bind* between a stored data version, its recorded dataset-level dependencies, and supplied actor metadata. The calling environment can provide prompt, run, agent-name, and script-path identifiers through R options or environment variables (e.g. `OL_PROMPT_ID`); when present, `put` /`save_as()` add them to commit metadata. If no identifiers are supplied, no agent context is inferred. `ol_log_commits` exposes stored versions, recorded parents, and available `agent_*` fields in process. An extensible adapter system brings Bioconductor containers into the same lake: the SummarizedExperiment adapter decomposes objects into assay, row/column, and metadata components for versioning and reconstruction, with adapters also available for SingleCellExperiment, MultiAssayExperiment, Spectra, MsExperiment, and QFeatures. Round-trip fidelity is assessed with class/structure checks and `all.equal(…, tolerance = 1e-8)` for numeric assays.

## 3 Results

We evaluated restoration and lineage capture under controlled conditions, fidelity on Bioconductor objects, use of supplied agent identifiers in a recovery case, and selected backend operation times.

### 3.1 Correctness of the versioning and lineage primitives

Across five reproducibility tests (RT-001–RT-005) based on ACM Artifact Review guidelines (ACM, 2020) (Supplementary Tables S1, S4 and S5), state restoration passed 30/30 independent iterations, each containing four nested state checks (120/120 checks; iteration-level 95% CI: 0.88–1.00). Numeric comparisons used `all.equal(…, tolerance = 1e-8)` after normalizing data-frame representation. In 60 generated pipelines restricted to tracked dplyr operations (427 derived tables and 619 ground-truth edges), automatic capture recovered all 619 edges without false-positive parents. This included 235 single-input edges and both parents of 192 joins (384 edges). After leaving tracked dplyr methods, however, capture was operation-dependent: an adversarial arm of 60 derivations achieved 40% recall, with three of five tested operation classes losing all edges. Such paths therefore require explicit dependencies. The remaining tests supported controlled intra-host transfer, version-count scaling within the prespecified threshold through 500 versions, and 5/5 rollback-cascade tests.

### 3.2 Fidelity on real Bioconductor omics

On the `airway` bulk RNA-seq dataset (Himes et al., 2014), a raw-count `SummarizedExperiment` round-tripped while being filtered and passed to DESeq2 (4,000 of 22,369 tested genes at adjusted *P <* 0.05). Explicit `depends_on` annotations recorded the filtering and DESeq2 boundaries, while the final dplyr join between differential-expression results and `rowRanges` -derived annotation captured both parents automatically; `tree()` recovered the resulting chain and the raw tagged state was restored. A tested 10x peripheral blood mononuclear cell (PBMC) `SingleCellExperiment` also preserved its sparse assay values and class, reduced dimensions, cluster labels, and time-travel state (Supplementary Table S6 and Figure S2). These cases demonstrate hybrid explicit and automatic lineage capture on the tested bulk and single-cell objects.

### 3.3 Recoverable agent provenance via the three-way bind

In a controlled, non-LLM case, we evaluated actor-associated provenance using 17,852 filtered airway genes (Supplementary Table S7). Environment variables supplied prompt, run, and agent identifiers for two fixed-threshold derivations: `runA` yielded 243 genes and `runB` yielded 4,505 genes. Both outputs retained their input lineage and supplied identifiers in commit metadata. In a matched overwrite experiment, runB replaced the live `deg` name, after which the runA commit was attributed through `ol_log_commits` and the runA intermediate was explicitly restored with its tag. The restored table matched the canonicalized reference values, columns, and row order. By contrast, the matched flat-file baseline overwrote `deg.csv` without a version or parameter record and could not attribute or recover the lost runA intermediate from its persisted state. A broader controlled suite detected all six injected provenance failures and executed restore actions for all four repairable auto-mode cases.

### 3.4 Backend operation benchmarks

Selected backend operations were benchmarked for 30 warm-cache iterations on an Apple M4 Max with 36 GB RAM (Supplementary Table S8 and Figure S3). Reading an existing 100 MB Parquet file with `arrow::read_parquet` took 0.0200 s versus 0.5716 s for `readRDS` (28.6 ×). With data preloaded in both backends, the tested DuckDB aggregation took 0.0062 s versus 0.0374 s for dplyr (6.0 ×), and the DuckDB join took 0.1631 s versus 0.6871 s for `base::merge` (4.2×). The join ratio is specific to that baseline: in a separate four-way comparison, data.table and dplyr were as fast as or faster than DuckDB. These measurements compare selected storage/query operations; they do not estimate the overhead of lineage capture itself.

## 4 Discussion

OmicsLake combines snapshots, tracked dplyr lineage, explicit dependencies, and optional actor metadata to support reproducibility of intermediate data states. In the four-tool comparison (Supplementary Table S3), OmicsLake was the only evaluated tool classified as providing all four tested capabilities: recoverable data versions, automatic ad hoc lineage, supplied actor identifiers, and in-process SQL. In the tested setup, `targets` exposed a predefined DAG but not recoverable history for overwritten target objects, while `pins` retained data versions without derivation lineage or agent fields. DuckLake was classified from its documentation as providing versioned tables and in-process SQL without automatic derived-from lineage or agent metadata; this comparison is not a systematic review of all provenance tools. The complete source-to-output map and tested boundaries are provided in Supplementary Tables S9 and S10.

The scope is deliberately limited. Automatic lineage applies to supported `ref()` –dplyr– `save_as()` paths; in-place mutation and operations crossing package boundaries require `depends_on` or a tracking context. Agent capture stamps identifiers supplied through options or environment variables and does not infer a human identity or execute an autonomous agent. The PBMC representation traded approximately 7.0× greater storage than `saveRDS` for queryable components and should not be interpreted as a compact sparse store. R memory limits also apply during conversion in `put()`, so datasets exceeding available memory require chunked processing.

Future work will extend tracking to additional Bioconductor methods, deepen integration with agent harnesses, and optimize sparse single-cell storage. OmicsLake is designed to support FAIR-aligned local workflows (Wilkinson et al., 2016) and complements Git, renv, and Docker across the code, environment, system, and intermediate data-state layers.

## Data and code availability

OmicsLake 0.99.3 is MIT-licensed at https://github.com/matsui-lab/OmicsLake (tag v0.99.3) (Matsui, 2026). The submitted source, tests, evaluation scripts, results, and per-analysis `sessionInfo` files are archived at https://doi.org/10.5281/zenodo.21391428. The evaluation used 0.99.1 (commit `1a3b81bef494`); subsequent releases do not alter the evaluated algorithms or reported values. The real-data case uses the public Bioconductor `airway` package (Himes et al., 2014).

## Acknowledgements

OpenAI Codex (GPT-5; July 2026) was used only to edit English. It did not generate or analyze data or determine conclusions; the author reviewed all changes and accepts responsibility. Details are in the Supplementary Material.

