## Supplemental Information for "OmicsLake: Versioned, Agent-Aware Data Lineage for R/Bioconductor Workflows"


Yusuke Matsui

Institute for Glyco-core Research (iGCORE), Nagoya University, Nagoya, Japan

Biomedical and Health Informatics Unit, Graduate School of Medicine, Nagoya University, Nagoya, Japan

ORCID: 0000-0003-3977-4313



Preprint version: 17 July 2026


**Scope of this supplement.** This document is organized around the claims evaluated in the accompanying preprint. It retains auditable protocols, additional results, authoritative source files, and scope boundaries while omitting repeated API exposition and redundant code. Generated CSV, text, and `sessionInfo` files are the numeric authority.

**Supplementary Table S1:** Claim-to-evidence map for the accompanying preprint.

| **Main-text claim** | Supporting evidence | Primary source |
|---|---|---|
| **Controlled restoration** | RT-001: 30/30 iterations and 120/120 checkpoints; 95% confidence interval (CI) 0.88–1.00. RT-005: 5/5 rollback cascades. | `02_reproducibility_test.R`; `results_reproducibility_summary.csv` |
| **Automatic lineage** | 60 randomized dplyr pipelines, 427 derived tables, and 619 true edges; recall and precision 100%. Adversarial non-dplyr recall 40%. | `10_lineage_accuracy_benchmark.R`; `results/lineage_accuracy_metrics.csv` |
| **Bioconductor fidelity** | Native airway `RangedSummarizedExperiment` and 10x peripheral blood mononuclear cell (PBMC) `SingleCellExperiment` round-trip, lineage, and time-travel checks. | `07_realdata_case_study.R`; `08_realdata_pbmc_case_study.R` |
| **Agent provenance** | Two controlled, fixed-threshold non-LLM runs produced 243 versus 4,505 genes; all four bind checks equaled 1; the overwritten runA state was attributable and explicitly recoverable from its tag. | `09_agent_provenance_case_study.R`; `11_recovery_baseline_contrast.R` |
| **Selected backend timings** | Warm-cache $n = 30$: existing-file Arrow read 28.6×, aggregation 6.0×, and join 4.2× versus selected baselines. No metadata-commit ratio is claimed because semantics differ. | `01_performance_benchmark.R`; `results/Table2A_core_benchmarks.csv` |
| **Comparative position** | OmicsLake, targets, and pins were measured in R; DuckLake was classified from published documentation. | `12_tool_comparison.R`; `results/tool_comparison.csv` |

### S1 Provenance contract and comparative scope

OmicsLake can associate stored versions with dataset-level, version-aware lineage, resolved parent versions, and supplied human or agent identifiers. It is an intermediate-state provenance layer, not a workflow orchestrator, and it does not infer every R operation.

**Critical capture boundary.** `lake$get()` → `lake$put()` does not create dependency edges. Automatic capture is limited to `ref()` → dplyr → `save_as()`; other paths require `depends_on` or `with_tracking()`.

**Supplementary Table S2:** Capture contract for common operation paths.

| Operation path | Capture | Required practice |
|---|---|---|
| `ref()` → filter/mutate/select/ summarize/arrange → `save_as()` | Automatic | Persist directly with `save_as()`; source metadata remains attached. |
| `ref(x)` → dplyr join(`ref(y)`) → `save_as()` | Automatic | Both x- and y-side parents are merged and recorded. |
| DESeq2, edgeR, limma, Seurat, scran/scater | Explicit | Use `put(..., depends_on=...)` or `with_tracking()`. |
| base/apply/transform/tidyr/custom or in-place operations | Explicit | Annotate each persisted result; do not assume automatic capture. |
| `collect()` followed by further transformation | Explicit | Lineage metadata may be lost after leaving the lazy tracked path. |
| `lake$get()` → arbitrary computation → `lake$put()` | Not automatic | Supply `depends_on` manually. |

#### Minimal tracked loop

The following schematic example shows the intended division between automatic dplyr capture and explicit annotation at a package boundary; `counts` is an input table and `dds` denotes a DESeq2 dataset constructed from the persisted `filtered` state by the surrounding analysis.

```
library(OmicsLake)
lake <- Lake$new("analysis")
options(ol.repro.capture = TRUE)

lake$put("counts", counts)
lake$ref("counts") |>
  dplyr::filter(total_count >= 10) |>
  save_as("filtered", lake)          # automatic parent: counts

fit <- DESeq2::DESeq(dds)
lake$put("de_results", as.data.frame(DESeq2::results(fit)),
         depends_on = "filtered")   # explicit package boundary
lake$snap("de_complete")
lake$tree("de_results")
```

#### Measured comparison

The comparison uses each tool's own metadata after a common minimal derivation. OmicsLake, targets, and pins were run in R. DuckLake was classified from published documentation because no CRAN R package was used in the experiment. "Partial" for targets denotes a predefined directed acyclic graph (DAG) and current-object cache rather than ad hoc lineage plus recoverable history.

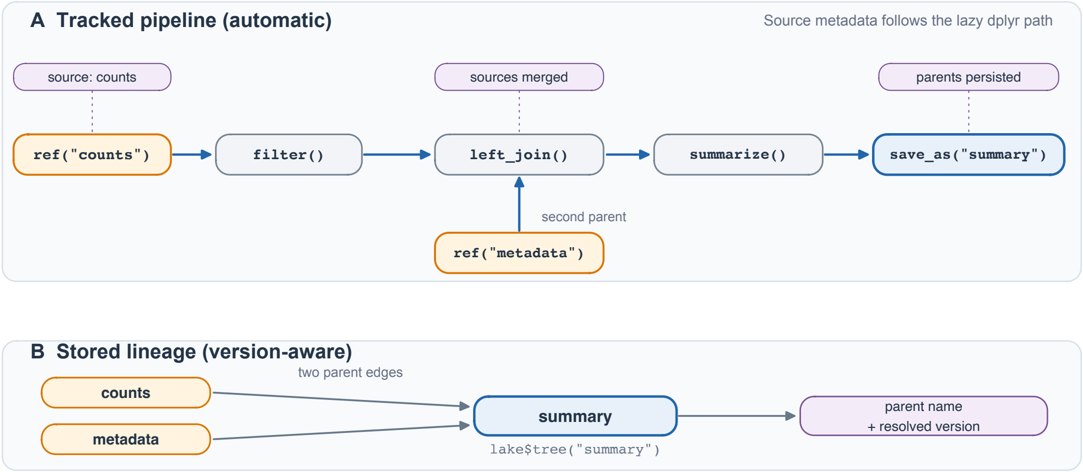


**Supplementary Figure S1:** Automatic lineage propagation through a tracked dplyr pipeline. Source attributes from `counts` and `metadata` are merged by the join and persisted by `save_as()`.

**Supplementary Table S3:** Head-to-head provenance capability comparison.

| Capability | OmicsLake | targets | pins | DuckLake |
|---|---|---|---|---|
| Recoverable data versions | Yes | Partial | Yes | Yes |
| Ad hoc automatic lineage | Yes | Partial | No | No |
| Supplied agent/actor identity | Yes | No | No | No |
| In-process query engine | Yes | No | No | Yes |

#### S2 Controlled validation of restoration and lineage

The validation separates integrity checks from benefit-level fault injection. Reproducibility tests RT-001 to RT-005 assess storage round-trip, lineage, same-host relocation, version accumulation, and rollback consistency. The breakage-taxonomy (BX) experiment injects realistic faults and tests detection, diagnosis, and recovery. The scaled lineage benchmark supersedes the original single nine-edge RT-002 graph as the headline lineage result.

**Supplementary Table S4:** Controlled reproducibility and recovery checks.

| Test | Scope | Design | Observed result | Interpretation |
|---|---|---|---|---|
| RT-001 | State restoration | 30 iterations; 4 checkpoints each | 120/120; 95% CI 0.88–1.00 | Storage-integrity foundation |
| Scaled lineage | Tracked dplyr DAGs | 60 pipelines; 619 true edges | Recall 100%; precision 100% | Headline automatic-capture result |
| RT-003 | Store relocation | Same-host `file.copy` only | 9/9 controlled checks passed | Not heterogeneous portability |
| RT-004 | Version accumulation | 50–500 versions | $< 2\times$ degradation | Bounded tested range |
| RT-005 | Rollback cascade | 5 project-state rollbacks | 5/5; 95% CI 0.48–1.00 | Small-$n$ supporting result |
| BX-001–006 | Injected failures | 6 fault classes | Detected 6/6; auto-restored 4/4 broken outputs | Benefit-level recovery evidence |

##### Scaled lineage-accuracy protocol

Arm A generated 60 seeded, valid DAG pipelines. Each derived table used either a tracked single-input dplyr verb or a two-input join and ended in `save_as()`; captured dependency edges were compared edge-for-edge with the recorded parent sets. Arm B routed 60 derivations through five non-dplyr operation classes (12 each) to measure the coverage boundary after leaving tracked methods.

**Supplementary Table S5:** Lineage-accuracy benchmark by derivation class. Recall and precision are micro-averaged over edges.

| Derivation class | Tables | Truth edges | Recall | Precision |
|---|---|---|---|---|
| Single-input dplyr verbs | 235 | 235 | 100% | 100% |
| Two-input dplyr joins | 192 | 384 | 100% | 100% |
| All dplyr-expressible | 427 | 619 | 100% | 100% |
| Adversarial non-dplyr | 60 | 60 | 40% | Not reported |

In the adversarial arm, apply-family, `base::transform`, and `tidyr::pivot_longer` lost all lineage in the tested implementations; `as.data.frame` plus base subsetting and the tested custom-function path retained lineage because they still resolved through a tracked path. Aggregate recall of 40% is therefore a measured property of this adversarial mixture, not a guarantee for arbitrary non-dplyr code. Such code should always be annotated explicitly.

##### Fault-injection interpretation

- BX-001 to BX-004 altered inputs, schema, parameters, or a critical artifact; each corrupted output was restored to the pre-fault hash.

- BX-005 and BX-006 removed Git or renv context. They were detected and produced guarded manual guidance, but no stored-data rollback was appropriate.
- The six scenarios constitute one run per scenario; they demonstrate mechanism coverage, not a population-level failure-rate estimate.

### S3 Bioconductor object fidelity on real data

Real-data studies test whether the storage and lineage primitives remain useful for native biological containers. Package-boundary transformations are intentionally recorded with `depends_on`; only the dplyr join between stored tables is automatically captured. This mixed strategy reflects the actual tracking contract rather than treating every biological operation as automatically observable.

**Supplementary Table S6:** Real-data object-fidelity and lineage checks.

| Case | Container and workflow | Measured results | Verified properties |
|---|---|---|---|
| airway bulk RNA-seq | `RangedSummarizedExperiment`; low-count filtering; DESeq2; annotation join | 63,677 raw genes; 22,369 tested; 4,000 with padj $< 0.05$; put 0.399 s; get 0.258 s | Structure/numeric round-trip; mixed manual+automatic lineage; raw-tag restoration |
| 10x PBMC single cell | `SingleCellExperiment` with sparse `dgCMatrix`; scran/scater clustering | 2,700 cells; 32,738 → 13,714 genes; 12 clusters; put 0.435 s | Sparse class/values; reduced dimensions; column labels; lineage; time-travel flags all passed |

#### airway provenance chain

The airway workflow stored the raw object, recorded a manual edge for Bioconductor filtering, recorded a second manual edge for DESeq2, derived gene annotation from `rowRanges`, and then joined DE results with annotation through the tracked dplyr path. `tree("de_annotated")` recovered both parents of the join and the upstream raw object. Restoring `@tag(raw)` recovered the original 63,677-gene state.

```
de_annotated <- lake$ref("de_results") |>
  dplyr::inner_join(lake$ref("gene_annotation"), by = "gene_id") |>
  save_as("de_annotated", lake)

# recovered chain:
# de_annotated <- de_results <- airway_filtered <- airway_counts
#              <- gene_annotation <- airway_counts
```

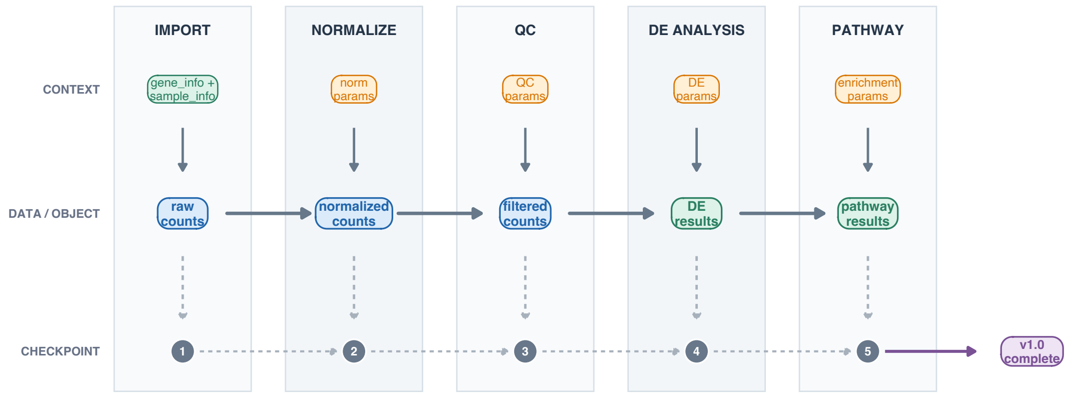


**Supplementary Figure S2:** Illustrative RNA-seq lineage DAG with typed data, object, parameter, commit, and version-label nodes. The measured airway chain is reported in the accompanying source text and result files.

##### Sparse-storage interpretation

The PBMC checks passed for sparse class and values, reduced dimensions, labels, lineage, and time travel. The on-disk footprint was 37.25 MB versus 5.35 MB for `saveRDS` (about 7.0× larger) because the current representation favors queryable long-format components over the compact original sparse object. This is an explicit storage trade-off, not a compression advantage.

#### S4 Recoverable agent provenance

**Scope.** This is a controlled test of provenance capture on real airway counts. It is not an autonomous large language model (LLM) execution and not a DESeq2 differential-expression test. Fixed fold-change thresholds stand in for two agent-produced alternatives.

Agent context was supplied through `OL_PROMPT_ID`, `OL_AGENT_RUN_ID`, and `OL_AGENT_NAME` under `options(ol.repro.capture=TRUE)`. OmicsLake passively stamped the resolved prompt, run, agent name, and script fingerprint onto each commit. Passive stamping supports attribution and audit; it does not orchestrate, sandbox, or validate the agent.

##### Controlled protocol and endpoints

1. Input: 17,852 low-count-filtered airway genes from the same stored data version.
2. Run A: `prompt-A-strong-DEGs` / `runA` / `analysis-agent`; $|\log_2 \mathrm{FC}| > 2$; 243 genes.
3. Run B: `prompt-B-lenient-DEGs` / `runB` / `analysis-agent`; $|\log_2 \mathrm{FC}| > 0.5$; 4,505 genes.
4. Verified flags: `three_way_bind_ok` and `lineage_ok`; `recovery_from_snapshot_ok` and `agent_attribution_ok`. All four equaled 1.

```r
Sys.setenv(OL_PROMPT_ID = prompt_id,
           OL_AGENT_RUN_ID = run_id,
           OL_AGENT_NAME = "analysis-agent")

lake$ref("airway_counts") |>
  dplyr::mutate(lfc = log2((trt_mean + 1)/(unt_mean + 1))) |>
  dplyr::filter(abs(lfc) > threshold) |>
  save_as(output_name)
```

**Supplementary Table S7:** Recovery contrast for the overwritten-intermediate scenario.

| Arm | Observed result | Why |
|---|---|---|
| With OmicsLake three-way bind | Attributable: yes; recoverable: yes; one explicit recovery action after attribution; canonicalized value/column/row-order hash match | The tagged version, bound commit metadata, lineage edge, and retained protocol context identify the runA state. |
| Conventional overwrite | Attributable: no; recoverable from persisted state: no; five attempted manual steps cannot recreate the missing state | `deg.csv` was overwritten; no prior version, parameter, lineage, or identity record remained. |

The claim is therefore narrower and more useful than "the agent is reproducible": OmicsLake makes an agent-associated intermediate attributable to a supplied run identity and makes a recorded state reversible. Reproducing the agent's reasoning, external services, or nondeterministic model output remains outside the tested scope.

### S5 Performance and storage interpretation

Benchmarks were run on one Apple M4 Max host with 36 GB memory, macOS 15.6 arm64, R 4.5.2, DuckDB 1.4.3, Arrow 22.0.0.1, dplyr 1.1.4, and data.table 1.18.0. Five warm-up iterations were excluded and the operating-system (OS) page cache was not cleared. Reported intervals are empirical 2.5th–97.5th percentile run intervals, not confidence intervals for a population mean.

**Supplementary Table S8:** Primary warm-cache performance measurements ($n = 30$). N/C denotes not comparable.

| Operation | Primary-method median [run interval] | Selected baseline | Ratio |
|---|---|---|---|
| Arrow read 100 MB | 0.0200 [0.0176–0.0407] s | `readRDS`: 0.5716 [0.5065–0.6529] s | 28.6× |
| Aggregate 1M rows | 0.0062 [0.0054–0.0074] s | dplyr: 0.0374 [0.0308–0.1448] s | 6.0× |
| Join 1M×1M | 0.1631 [0.1365–0.2085] s | `base::merge`: 0.6871 [0.5518–1.1209] s | 4.2× |
| Metadata commit | 0.1255 [0.1096–0.2429] s | `file.copy`: 0.0073 [0.0040–0.0107] s | N/C |

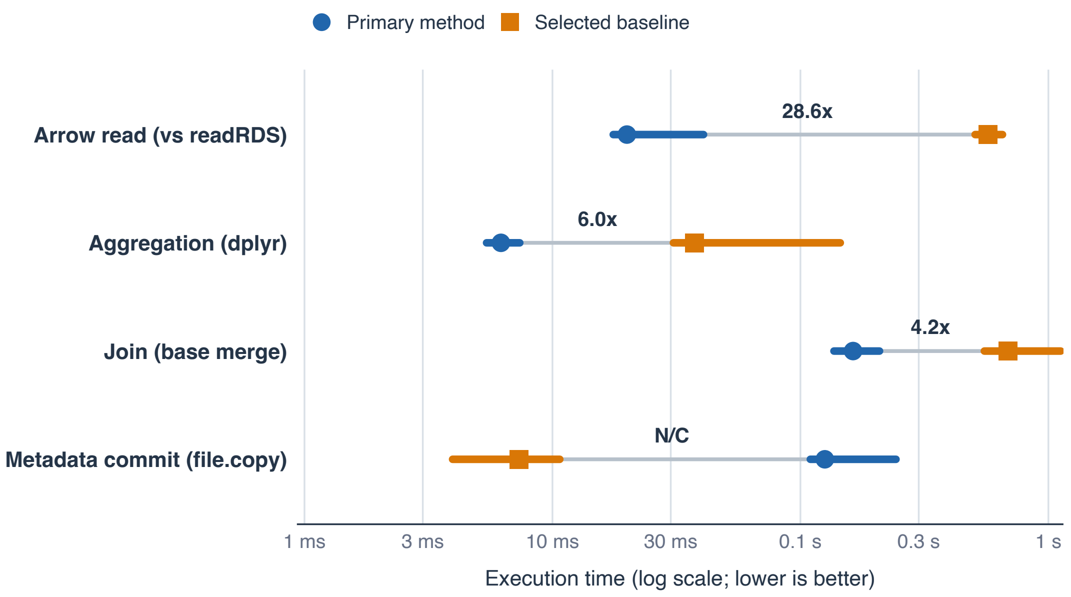


**Supplementary Figure S3:** Warm-cache benchmark medians and empirical run intervals. Metadata commit and file copy are shown without a speedup ratio because they do not implement the same semantics.

**Interpretation guardrails.** The import row times `arrow::read_parquet()` versus `readRDS()` on existing files; it does not include DuckDB registration or a complete OmicsLake ingest. The aggregation and join rows time selected calls after data were stored or materialized, and no benchmark compares lineage capture enabled versus disabled. The 4.2× join ratio is versus `base::merge`; in a separate four-way block, data.table (0.0551 s) and dplyr (0.1382 s) were as fast as or faster than DuckDB (0.1439 s). These are selected backend-operation measurements, not universal in-memory superiority or a provenance-overhead estimate.

##### Storage trade-offs

- Dense numeric benchmark: Parquet 35.2 MB versus compressed RDS 29.5 MB (19.3% larger).
- airway `RangedSummarizedExperiment`: OmicsLake 42.37 MB versus `saveRDS` 9.88 MB because of queryable long-format decomposition.
- PBMC sparse `SingleCellExperiment`: OmicsLake 37.25 MB versus `saveRDS` 5.35 MB (about 7.0× larger).
- OmicsLake is therefore presented as queryable, versioned provenance storage—not as the smallest representation for every omics object.

#### S6 Reproduction map and limitations

Generated CSV, text, and `sessionInfo` files are the source of quantitative values in this supplement. The concise map below identifies each script and its direct outputs; narrative summaries are not treated as the numeric authority.

**Supplementary Table S9:** Reproduction map from experiment scripts to authoritative outputs.

| Evidence module | Run script | Authoritative outputs |
|---|---|---|
| Performance and storage | `inst/paper/01_performance_benchmark.R` | `results/Table2A_core_benchmarks.csv`; `results/Table2B_storage_efficiency.csv`; `results/Table2C_fair_comparison.csv` |
| RT-001–RT-005 | `inst/paper/02_reproducibility_test.R` | `results_reproducibility_summary.csv`; `results_rt004_scalability.csv` |
| Breakage taxonomy | `inst/paper/04_breakage_taxonomy_validation.R` | `results_breakage_evaluation.csv`; `results_breakage_coverage_summary.csv` |
| Lineage accuracy | `inst/paper/10_lineage_accuracy_benchmark.R` | `results/lineage_accuracy_metrics.csv`; `results/lineage_accuracy_adversarial_by_op.csv` |
| airway case | `inst/paper/07_realdata_case_study.R` | `results/realdata_metrics.csv`; `results/realdata_lineage_trees.txt`; `results/realdata_sessionInfo.txt` |
| PBMC case | `inst/paper/08_realdata_pbmc_case_study.R` | `results/pbmc_metrics.csv`; `results/pbmc_lineage_tree.txt`; `results/pbmc_sessionInfo.txt` |
| Agent bind | `inst/paper/09_agent_provenance_case_study.R`; `inst/paper/11_recovery_baseline_contrast.R` | `results/agent_provenance_metrics.csv`; `results/recovery_contrast_metrics.csv`; `results/agent_provenance_sessionInfo.txt` |
| Tool comparison | `inst/paper/12_tool_comparison.R` | `results/tool_comparison.csv`; `results/tool_comparison_sessionInfo.txt` |

#### Consolidated limitations

**Supplementary Table S10:** Tested boundaries and appropriate interpretation.

| Boundary | What was tested | What is not established |
|---|---|---|
| Lineage granularity | Dataset-level parents with resolved version references | Column-, cell-, or operation-level provenance |
| Automatic capture | Tracked `ref()` → dplyr → `save_as()`, including joins | Arbitrary R/Bioconductor operations without annotation |
| Agent support | Passive prompt/run/agent stamping and recorded-state recovery | Agent reasoning, orchestration, sandboxing, or validation |
| Portability | Same-host copied-store checks | Heterogeneous architecture/runtime portability |
| Sparse storage | Correct class/value round-trip on PBMC | Compact sparse-native storage at large scale |
| Performance | Warm-cache single-host selected comparisons | Universal speed advantage, cold-cache behavior, or multi-host scaling |
| Study size | 30 restoration iterations; 60 lineage DAGs; one run per BX scenario | Population failure rates or broad platform generalization |

### S7 AI-assisted language editing disclosure

During manuscript preparation, the author used OpenAI Codex (GPT-5; accessed July 2026) solely for language-level revision of author-prepared manuscript passages, including improvements to English grammar, clarity, and concision. The tool was not used to generate or analyze research data, select results, produce figures, or determine scientific interpretations or conclusions. The author critically reviewed and revised every accepted suggestion and takes full responsibility for the final manuscript.

End of Supplementary Materials
